# Modelling the contribution of iodised salt in industrially processed foods to iodine intake in Macedonia

**DOI:** 10.1101/2020.12.20.423637

**Authors:** Neda Milevska-Kostova, Borislav Karanfilski, Jacky Knowles, Karen Codling, John H Lazarus

## Abstract

Evidence from the 1950s showed that Macedonia was iodine deficient. After the introduction of mandatory universal salt iodisation, the country saw a steady increase in iodine intake and decline in goitre prevalence, earning iodine-deficiency free status in 2003. Iodine status assessments in 2007 and 2016 showed adequate iodine intake among school age children (median urinary iodine concentration of 241 µg/L and 236 µg/L respectively). Macedonia participated in the 2019 piloting of the Iodine Global Network Programme Guidance on the use of iodised salt in industrially processed foods to better understand salt and iodised salt intake from food sources other than household salt.

Aggregated data from the 2017 Household Consumption and Expenditure Survey (HCES) was used to determine household salt consumption, to identify widely-consumed, salt-containing industrially processed foods and estimate typical daily intake of these foods. The salt content of these foods was estimated using national standards and the Danish food composition database. The percentage of this salt that was iodised was assessed using customs data for salt imports.

Although the study has its limitations, including a relatively small selection of foods, the results indicate potential iodine intake from iodised household salt and iodised salt in the selected foods of above 300% of the Estimated Average Requirement and over 220% of the Recommended Nutrient Intake for adults. This was approximately 50% of the tolerable safe Upper Level for iodine intake. The study confirmed high daily salt intake (11.2 grams from household salt only). Successful salt reduction would be expected to reduce iodine intake, however, modelling with 10% and 30% reduction implied this is unlikely to put any population group at risk of deficiency. It is recommended that design and implementation of salt iodisation and salt reduction policies are harmonized, alongside continued regular iodine status monitoring for different population groups.

## Introduction

There is evidence that Macedonia, a land-locked country in Southern Europe, was historically iodine deficient, with goitre prevalence of up to 30% in some regions [1, 2]. According to a study by Ramzin, in 1953 Macedonia had nearly 200,000 inhabitants (or more than 20% of total population) with goitre [3]. To improve iodine intake, mandatory iodisation of salt for human and animal use with 10 mg iodine (as potassium iodide) per kg salt was introduced in the former Yugoslavia in 1956 [4]. Decline in goitre prevalence was recorded in 1980s [5], however, the first nationwide iodine status assessment was conducted post-independence in 1996. This survey showed median urinary iodine concentration (MUIC) of 117 µg/L among 2,380 schoolchildren and goitre prevalence of 18.7% among the 11,486 schoolchildren tested [3]. To sustain adequate iodine intake among children and further reduce the goitre prevalence, the Macedonian government established a multisectoral Committee on Iodine Deficiency in 1997 and enacted a new national policy on mandatory iodisation of all food grade salt in 1999, with 20-30 mg of iodine per kg salt, using potassium iodate which is more stable than the iodide form. Regular assessments of iodine status in schoolchildren have shown a steady increase in iodine intake, reaching MUIC of 241 µg/L in 2007, and steady decline in goitre prevalence to <1% in 2007 [1]. The 2016 iodine status re-evaluation confirmed proper implementation of the salt iodisation program, suggesting optimal iodine intake among schoolchildren (MUIC of 236 µg/L) [5].

At the same time, literature shows that the population in Macedonia has very high salt intake, estimated at 14 grams per person per day [6]. High salt consumption affects blood pressure, and high blood pressure is key risk for cardiovascular diseases. Sodium consumption of more than 2g per day is estimated to cause 1.65 million cardiovascular related deaths each year, or 1 of every 10 deaths from cardiovascular causes [7]. Salt reduction was listed by the 2011 UN high-level meeting on non-communicable diseases, as one of the top three priority actions to reduce premature mortality from non-communicable diseases [8]. The World Health Organisation in its 2013 guideline recommends a 30% reduction in sodium intake by 2025, with an ultimate goal of reaching daily consumption of 51g salt (2g sodium) for adults worldwide [9]. To achieve this target, Macedonia needs to introduce a salt reduction policy, that should be informed by evidence on dietary habits and main sources of salt. Reducing salt consumption would reduce iodine intake, however, the extent of this effect is difficult to quantify without understanding the contribution of, potentially iodised, salt in industrially processed foods to salt and iodine intake.

Our national team submitted an expression of interest to pilot the draft Iodine Global Network (IGN) Programme Guidance (IGN PG) on the use of iodised salt in industrially processed foods in 2019. The aim was to improve understanding of national dietary habits and the potential and estimated current iodine intake from iodised household salt and from iodised salt in widely consumed industrially processed foods. One of the intended objectives from implementing the IGN PG was to identify the need, opportunities, and required actions to strengthen the processed food component of the national salt iodisation policy.

## Materials and methods

A national working group comprised of representatives from public health and clinical community, government, consumer association and industry representatives was established to provide feedback and validation of results of the piloting. The assessment was led by two national focal points with a combined background in endocrinology, iodine nutrition, research methods and policy analysis. Implementation took place between March and December 2019 following the IGN PG framework, described by Knowles et al (in-print). To ensure coherence of the process, online consultative meetings with IGN technical advisors took place upon completion of each module. Findings and results were compiled in an assessment report, that served as basis for development of a draft-action plan. The assessment report and draft-action plan were shared with the national working group for feedback and validation.

An initial step of the assessment was to identify potential data sources for salt containing processed food consumption, salt intake, processed food salt content and the percent of household salt and food industry salt that was iodised. The 2017 data from the Household Consumption and Expenditure Survey (HCES) were used to identify widely consumed processed food products that potentially represent the most significant sources of salt. As individual level consumption data were not possible to obtain from the Macedonian State Statistical Office (SSO), we used aggregated data publicly available from the SSO website [10]. The HCES reports included information on annual consumption of individual foods per household member, not calculated as adult male equivalence values. The average daily per capita consumption of each selected food was determined by dividing the average annual consumption per household member by number of days in a calendar year (365 days).

To estimate the salt content in these foods, we referred to the available normative standards in the country; namely, we used the bylaws of the Law on food safety [11] that regulate the maximum allowed salt content of milk and dairy products [12], and of soups and sauce concentrates [13]. For foods for which no normative for salt content was found in the bylaws (bread, sausage, salami and ham), we used the indicated average sodium content from the Danish database of food composition tables [14]. To convert sodium to salt content, we used the conversion formula (m(NaCl) = m(Na) × 2.54).

As the entire salt in the country is imported, we obtained data from the Customs authority on salt imports by quantity and importer for 2014 to 2018, to determine the percentage of iodised salt used. Initially, data was filtered to include only importers of food grade salt (household and food industry), and then analysed to estimate the relative percentage of iodised and non-iodised salt imported in total.

We then used the IGN PG Excel-based tool to model the potential contribution of typical consumption patterns for the identified processed foods to each of the: estimated average requirement (EAR) for iodine, the recommended nutrient intake (RNI) for iodine, and the tolerable upper limit (UL) for iodine intake [15]. The potential iodine intake from consumption of household salt and each processed food was based on an assumption that 100% of food grade salt was iodised to the mean of the national standard for Macedonia (20 to 30 mg/kg) and accounted for up to 30% loss of iodine in the final product at time of consumption. The assessment was run again using estimates for estimated current percent iodised salt used by households (100%) and in the food industry (94%).

As successful implementation of a future national salt reduction policy would lead to reduced iodine intake from salt sources, we attempted to assess the potential impact of salt reduction on the availability of iodine under the current iodisation levels, and whether effective implementation of salt reduction policy would require adjustments to the salt iodine standard. We used the same IGN PG Excel-based tool to model the potential influence of an effective salt reduction policy on iodine intake, in order to obtain an idea of the expected change in iodine intake from household salt and from iodised salt in these processed foods if national salt reduction targets of 10% or 30% were achieved. The two targets for modelling were chosen based on: 1) the experience of other countries (i.e. 10% reduction based on a 2% annual reduction until 2025), and 2) WHO recommendations (i.e. 30% reduction) [9].

In the following step, the national focal points conducted a review of enabling factors and any gaps in these, related to the existing strategy and policy to achieve optimal iodine intake.

The outcomes of the modelling and broader programme review were presented to the National Committee on Iodine Deficiency, established by the Ministry of Health with an advisory role in monitoring and development of evidence-based recommendations.

## Results

Based on the identified available data, we conducted the assessment of potential contribution to iodine intake from the use of iodised household salt and of iodised in selected processed foods, shown in Figure 1 of Knowles et al (in-print), based on typical consumption patterns for the identified foods.

**Fig 1.**
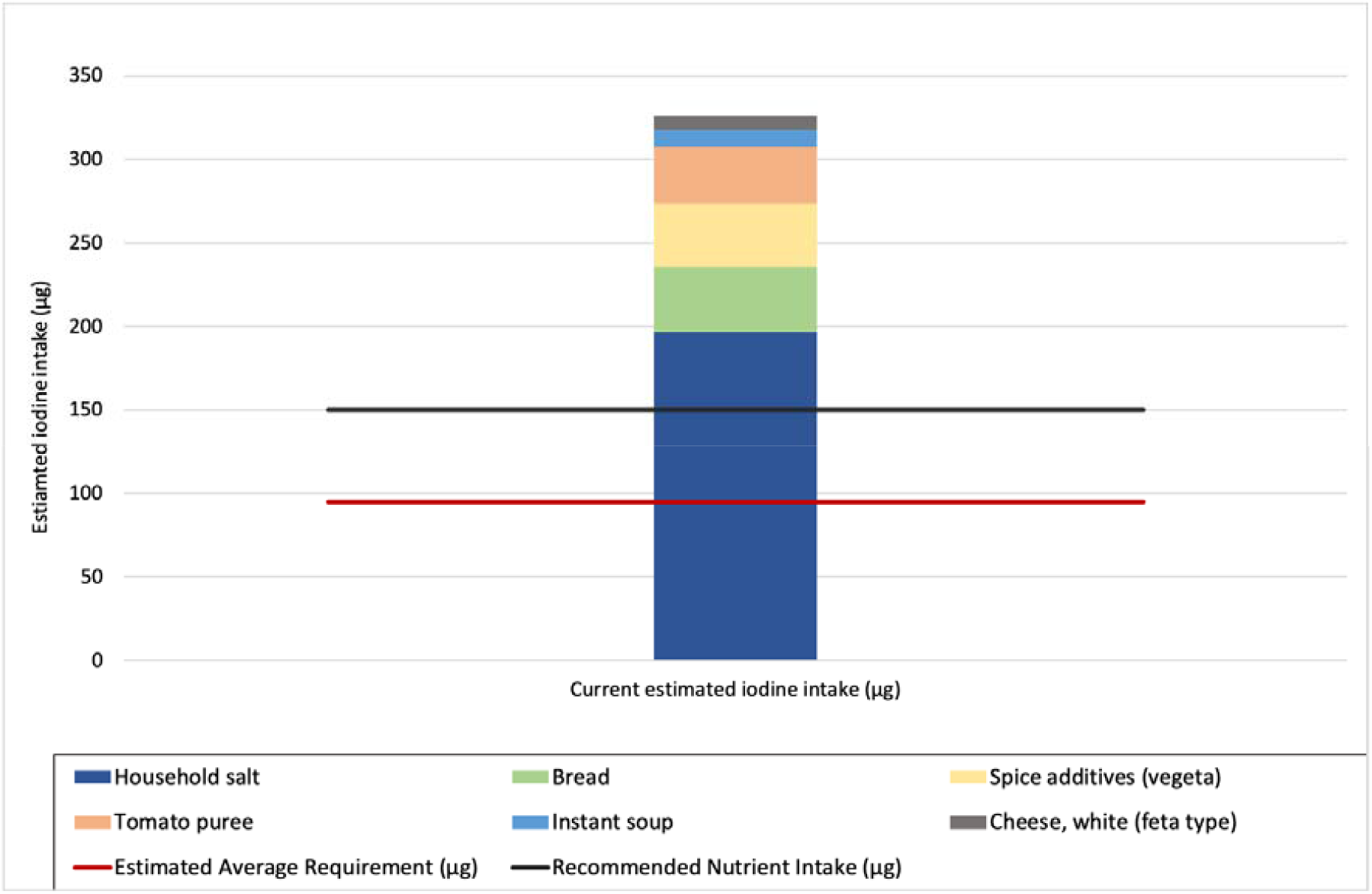
Contribution of household salt and iodised salt in selected processed foods to estimated current iodine intake, in relation to the EAR and RNI for iodine. The values for iodine intake are based on knowledge of current iodisation practices (100% household salt and 94% food industry salt iodised). Note that the tolerable upper level (UL) for iodine intake of 600 µg iodine is not shown on this chart.

### Selection of industrially processed foods for modelling

The process for selection of relevant salt-containing processed food products from HCES for the modelling exercise had several steps. Firstly, we excluded 85 products for which the relative error of annual household intake as reported by SSO was >10%. Of the remaining 67 foods, we further filtered the list to contain only household salt and industrially produced foods (n=29). Within the processed foods, we identified foods that either have: i) low to medium salt content but high overall consumption (e.g. bread) or ii) high salt content and consumed frequently, even if in low amounts, that can still significantly contribute in the overall salt intake. The final list of industrially processed foods used in the modelling exercise included nine products: household salt, bread, spice mix (vegeta), tomato puree, instant soup, white cheese white (feta type), sausages (various), ham and salami. We then looked at the availability of national standard for salt content in each of these products, and further eliminated products for which: 1) there was no national standard; and/or 2) the wide variety of types classified under that product made it difficult to estimate the salt content (e.g. sausages, ham and salami). Although there was no national standard for it, bread was retained for analysis due to the high daily consumption of this product. The salt content of bread was estimated using the Danish food composition database.

Table 1 shows the average daily per capita consumption and estimated salt content of household salt and the processed foods selected for modelling (n=6).

**Table 1.** Average daily consumption and salt content of industrially processed foods selected for modelling.

| Food product | Estimated average daily per capita consumption (g) | Salt content (% product weight) |
| --- | --- | --- |
| Household salt | 11 | 100.0% |
| Bread | 198 | 1.2% [14] |
| Spice mix (vegeta) | 4 | 60.0% [13] |
| Tomato puree<sup>a</sup> | 8 | 25.0% [13] |
| Instant soup | 1 | 45.0% [13] |
| Cheese, white (feta type) | 16 | 3.0% [12] |

<sup>a</sup> salt standard for all sauces

### Determining the percent of food grade salt that is iodised

To estimate the percentage of iodised salt used in the country we first looked at the salt import inspection policies and practices. While monitoring is well-established for household salt, it presents somewhat of a challenge when it comes to salt used in food industry. The entire salt used in the country is imported and its iodisation levels are subject to inspection at the point of entry into the country.

According to the Macedonian Customs authority data for 2014 to 2018, there are two large salt importers covering between 97.7% (in 2014) and 94.9% (in 2018) of all salt imports in the country; between 75 to 85% of food grade salt is imported by the largest importing company, followed by 12 to 16% imported by the next largest importer. The remaining 4 to 5% is imported by various small companies. Due to their market share size, the Food and Veterinary Agency performs regular inspection controls in the two largest importing companies. Such finding suggests that more or less all salt used in both households and the food processing industry in the country can be assumed to be iodised.

### Determining average daily salt and iodine consumption

Based on HCES 2017 data, the average consumption of household salt per household member was 11.2 grams per day. Adding the salt consumed through the industrially processed foods selected for modelling, further increased the average daily salt consumption. Estimated current iodine intake through household salt and salt used in the chosen products was based on an estimate that 100% of household salt and 94% of food industry salt is iodised to the mean of national standards (20 to 30 mg/kg). Using these consumption and salt iodisation estimates, iodine intake from household salt alone is above both the RNI and EAR for iodine. Adding intake from iodised salt in the selected processed foods provides a total estimated daily intake of 325 µg iodine, which is approximately 330% of the EAR and 210% of the RNI for iodine for non-pregnant adults (Fig. 1). These levels indicate adequate iodine intake that remains well below the tolerable UL of iodine intake of 600µg for this population group.

### Modelling the potential impact of a salt reduction policy on iodine intake

Due to the established association between high salt consumption and cardiovascular diseases [7], the World Health Organisation recommends a 30% reduction by 2025, with an ultimate goal of reaching daily consumption of 5⍰g salt for adults worldwide [9].

To achieve this target, Macedonian authorities would have to introduce and implement a very stringent and assertive salt reduction regimen. We have modelled the potential impact of salt reduction policy on iodine intake, using two distinct targets, as described in the methods section, i.e. 10% and 30% reduction.

Fig. 2 shows the outcome of modelling the potential impact of salt reduction on iodine intake from the selected foods where the target is 10% and 30% reduction in all foods.

**Fig. 2.**
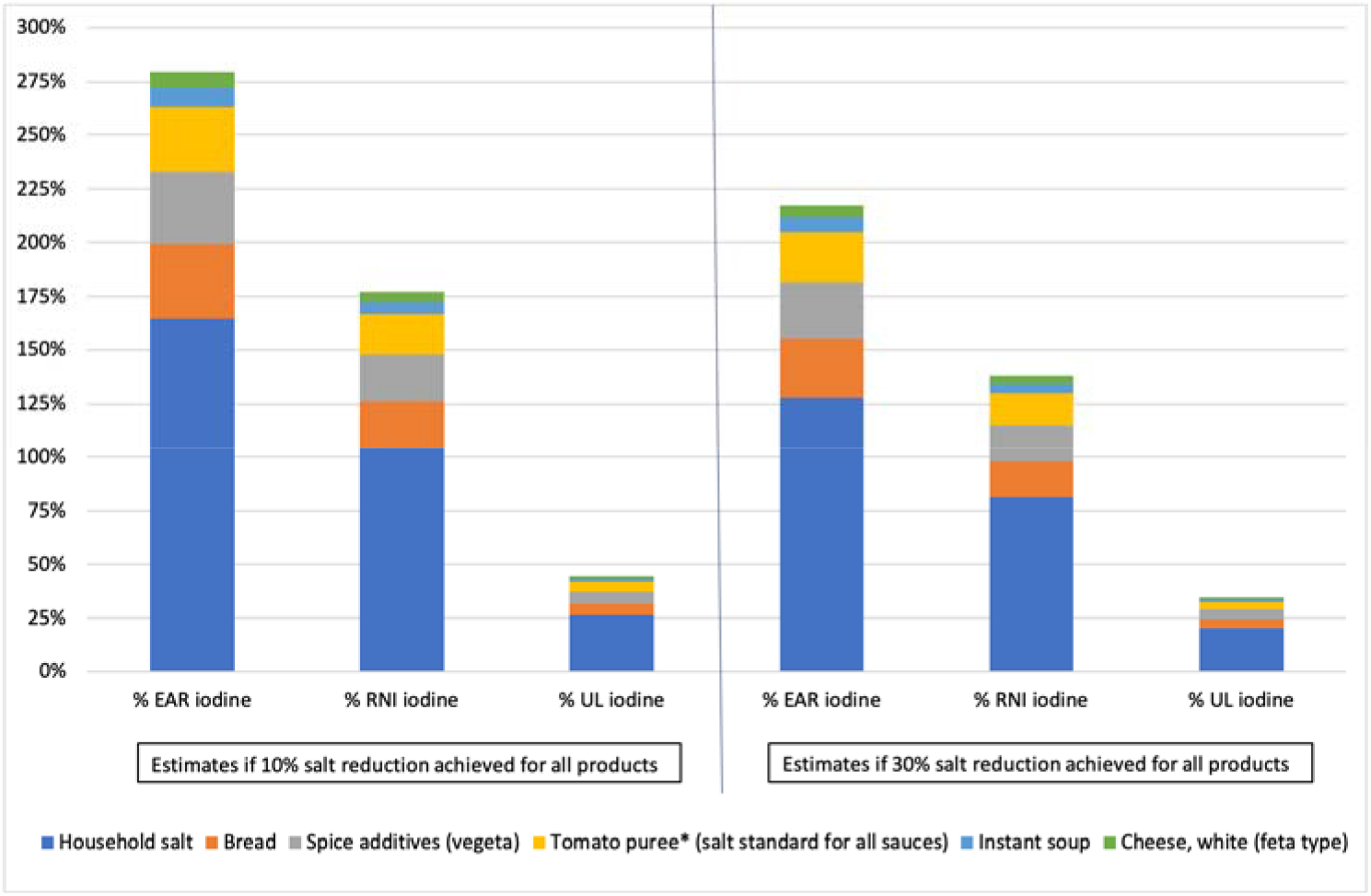
Potential impact of successful salt reduction on iodine intake from household salt and processed food salt (10% and 30% reduction scenarios) Potential (100% household salt iodised and 94% industry salt iodised) estimated % EAR iodine, % RNI iodine and % UL iodine before and after successful salt reduction

If 10% salt reduction was achieved in consumption of household salt and in consumption of salt from the selected processed foods and salt iodine levels and percent of salt iodised remained the same, then the contribution to % nutrient reference values for iodine from household salt and salt in the selected products would reduce from 330% to 280% of the EAR, from 210% to 180% of the RNI, and from 55% to 45% of the UL (Fig. 2).

Under the same conditions described above, if 30% salt reduction is achieved, then the contribution to the % nutrient reference values for iodine from household salt and salt in the selected products would reduce to 220% of the EAR, to 140% of the RNI, and to 35% of the UL (Fig. 2).

### Limitations

Our study had several limitations. Firstly, the level of data aggregation available from the HCES dataset resulted in approximations in estimating total daily salt intake per household member. To reduce the effect of this limitation, we only included food products for which data were reported with <10% error for household level intake. A second limitation was the availability of information regarding the salt content for the selected processed foods. We used national normative standards where these were available, and a non-specific source (the Danish food composition database) for the remaining products. To minimise the impact of this limitation, we excluded processed foods where a large range of salt content was found for slightly different varieties of the same product, such as the processed meat products (sausages, salami and ham). Finally, the study could not factor in the salt and iodine content of industrially processed foods that were imported. As explained in the methodology section, the calculations were based on the assumption that 94% of selected processed foods were produced using salt iodised to the mean of the national standard. However, for the period for which data were examined (2017), the country’s total imports in the food category surpassed exports by 26.6% [16]. This allows for the conclusion that some portion of the consumed food is likely to have been imported from other countries and these countries may have different normative standards for product salt content and salt iodisation. The impact of this potential limitation was reduced by the fact that most bread, tomato paste and white cheese would have been locally produced. Bearing in mind that most effective salt reduction strategies include specific product reformulation, an additional limitation is the application of the same % salt reduction across all products.

## Discussion

Continuous provision of sufficient iodine in human nutrition is key to successful prevention of iodine deficiency, and universal salt iodisation (USI) is one of the most cost-effective measures to achieve this [17]. Based on regular iodine status evaluation, it can be concluded that iodine status of all population groups in Macedonia is sufficient[1, 5, 18], which still represents a challenge in many countries [19, 20]. This also implies that the current level of iodine in salt (20 to 30 mg/kg salt) is appropriate.

The results of the modelling exercise to determine contribution of iodised household salt and iodised salt in industrially processed foods to iodine intake of the general population confirm that sufficient iodine intake to meet population requirements is possible from these products. However, once achieved, the iodine deficiency improvement is not a permanent benefit. Reduction of both attention and activities that contribute to ensuring proper iodisation of all food grade salt might result in inadequate iodine intake and bring back all related consequences of iodine deficiency. Thus, the WHO, UNICEF and IGN in many of their documents emphasize the importance of achieving and sustaining USI, and recommend regular and compulsory re-evaluation of iodine status and of USI even in countries with achieved iodine sufficiency, both to not fall back into deficiency, as well as to avoid moving to the other extreme of above optimal iodine and its negative health consequences. Based on the findings in this study, as well as of the assessment of iodine intake of school children conducted in 2016, Macedonia should continue with regular monitoring of both iodised salt quality and iodine intake. In addition, iodine assessment of pregnant women has been sporadically conducted since 2007 [1, 21] and this should be continued, whereas other groups, such as nursing mothers and infants were so far not considered for a nation-wide study.

However, while the findings for iodine intake in this study reassures of good prevention of iodine deficiency disorders, the very high intake of salt presents a case for concern, implying the need for introduction of salt reduction policy in the country. Due to the established association between high salt consumption and cardiovascular diseases [7], the World Health Organisation recommends a 30% reduction by 2025, with an ultimate goal of reaching daily consumption of 5⍰g salt for adults worldwide [9]. To achieve this target, Macedonian authorities would have to introduce and implement a very stringent and assertive salt reduction regimen. Literature shows that even the most organized multi-faceted approaches require long-term commitment and take a long time to yield results [6, 22]. For example, in the United Kingdom, a 7-year period of implementing salt reduction strategies contributed to 2% annual reduction of salt intake [23].

One approach to salt reduction is to introduce food-specific reformulations with adjustments to salt content. For this, processed foods are selected based on an assessment of their contribution to salt intake among different population groups and consumer acceptability of a reduced salt product. Evidence for the initial identification of potential processed foods can be drawn from a larger scale study, that can use our assessment as a starting point.

## Conclusion

Even though the assessment presented in this paper provides only an indication of the estimated total salt and iodine intake through household salt and some processed food products, it can be concluded that the general population has sufficient access to iodine as a result of a well-implemented universal salt iodisation policy. However, the results from this assessment indicate generally high population salt intake which highlights a need for additional research to further determine total salt intake and its main sources, to develop an effective policy for salt reduction. A salt reduction policy could target individual level consumption (for example, behaviour change to reduce the use of household salt and to consume foods with lower salt content), and or the food industry (for example, product reformulation discussed above).

Results of the study reassure that achieving even the most optimistic salt reduction scenario (30% reduction) should not lead to inadequate iodine intake. Therefore, there will likely be no need to make any revisions to the current salt iodisation standards. However, to be assured of sustained optimal iodine intake, it is recommended that salt iodisation and salt reduction policies are designed, implemented and monitored in a coordinated manner [6]. Regular iodine status assessment practice should be continued for both general and specific vulnerable population groups, such as pregnant and nursing women and infants, alongside continuous regulatory monitoring and enforcement of food grade salt iodisation.

## Acknowledgements

The authors would like to thank Prof. Dr Daniela Miladinova and Assoc. Prof. Dr Igor Spiroski, Faculty of Medicine, University “Ss. Cyril and Methodius”, Marijana Loncar Velkova, PhD, President of the Consumers’ Organization of Macedonia (COM), Mr. Goran Angelovski, Director of the Salt iodisation factory “Izvor” and Member of the National Committee for Iodine Deficiency, Ministry of Health and Mr. Miroslav Balaburski, Food and Veterinary Agency for their contribution as part of the national working group.

## References

1. Karanfilski B, Milevska-Kostova N, Miladinova D, Jovanovska V, Kocova M. Continued efforts are key to sustaining iodine sufficiency in Macedonia. IDD Newsletter. 2018;4.

2. Karanfilski B, Sestakov G, Loparska S, Miceva-Ristevska S, Serafimov N, Tadzer I, et al. Reevaluation of the results of iodine prophylaxis in eradication of endemic goiter in Macedonia. Med Pregl. 1993;46 Suppl 1:77–9. PubMed PMID: 8569616.

3. Karanfilski B, Bogdanova V, Vaskova O, Loparska S, Miceva-Ristevska S, Sestakov G, et al. Correction of iodine deficiency in Macedonia. J Pediatr Endocrinol Metab. 2003;16(7):1041–5. PubMed PMID: 14513882.

4. Buzina R. Ten years of goiter prophylaxis in Croatia, Yugoslavia. The American journal of clinical nutrition. 1970;23(8):1085–9.

5. Majstorov V, Miladinova D, Kuzmanovska S, Ittermann T, Gjorcheva DP, Vaskova O, et al. Schoolchildren thyroid volume in North Macedonia: data from a national survey in an iodine-sufficient country. Journal of Endocrinological Investigation. 2020:1–7.

6. Trieu K, Neal B, Hawkes C, Dunford E, Campbell N, Rodriguez-Fernandez R, et al. Salt reduction initiatives around the world–a systematic review of progress towards the global target. PloS one. 2015;10(7):e0130247.

7. Mozaffarian D, Fahimi S, Singh GM, Micha R, Khatibzadeh S, Engell RE, et al. Global sodium consumption and death from cardiovascular causes. New England Journal of Medicine. 2014;371(7):624–34.

8. Beaglehole R, Bonita R, Horton R, Adams C, Alleyne G, Asaria P, et al. Priority actions for the non-communicable disease crisis. The Lancet. 2011;377(9775):1438–47.

9. World Health Organization. WHO issues new guidance on dietary salt and potassium. WHO Media Centre. 2013.

10. SSO. Household consumption in the Republic of Macedonia, 2017. Skopje: State Statistical Office, 2018.

11. Law on food safety, Official Gazette of RM, no. 157/2010, (2010).

12. Rulebook on the special safety and hygiene conditions and way of performing controls of milk and dairy products, Official Gazette of RM, no. 26/2012, (2012).

13. Rulebook on requirements on quality of soups, soup concentrates, sauce concentrates, spice additives and similar products, Food and Veterinary Agency, Official Gazette of RM, no. 95/2012, (2012).

14. Christensen T, Biltoft-Jensen AP, editors. The New version of Danish food composition database FRIDA including a case study on recipe calculation compared to a chemical analysis. 39th National Nutrient Databank Conference; 2016.

15. Allen LH, Carriquiry AL, Murphy SP. Perspective: proposed harmonized nutrient reference values for populations. Advances in Nutrition. 2020;11(3):469–83.

16. External trade by enterprise characteristics, 2017 [Internet]. State Statistical Office; 2019. Available from: http://www.stat.gov.mk/pdf/2019/7.1.19.08_mk.pdf

17. WHO. Guideline: fortification of food-grade salt with iodine for the prevention and control of iodine deficiency disorders. 2014.

18. Karanfilski Be. Iodine and thyroid status of the population in Macedonia, 2018 (monograph). Skopje: Medical Faculty, University “Ss. Cyril and Methodius”; 2018. 199 p.

19. Dold S, Zimmermann MB, Jukic T, Kusic Z, Jia Q, Sang Z, et al. Universal salt iodization provides sufficient dietary iodine to achieve adequate iodine nutrition during the first 1000 days: a cross-sectional multicenter study. The Journal of nutrition. 2018;148(4):587–98.

20. IGN. Global Scorecard of Iodine Nutrition in 2017 in the general population and in pregnant women (PW). IGN: Zurich, Switzerland. 2017.

21. Kuzmanovska S, Majstorov V, Miladinova D, Jovanovska V, Atanasova Boshku A, Shabani A, et al. Iodine supplementation and thyroid status in healthy pregnant women in iodine-replete region. Macedonian Medical Preview. 2019.

22. Webster JL, Dunford EK, Hawkes C, Neal BC. Salt reduction initiatives around the world. Journal of hypertension. 2011;29(6):1043–50.

23. He F, Brinsden H, MacGregor G. Salt reduction in the United Kingdom: a successful experiment in public health. Journal of human hypertension. 2014;28(6):345–52.

